# Mitochondrial sirtuins *sir*-*2.2* and *sir*-*2.3* regulate lifespan in C. *elegans*

**DOI:** 10.1101/181727

**Authors:** Sarah M. Chang, Melanie R. McReynolds, Wendy Hanna-Rose

## Abstract

Mitochondrial sirtuins regulate biochemical pathways and are emerging drug targets for metabolic and age-related diseases such as cancer, diabetes, and neurodegeneration. Yet, their functions remain unclear. Here, we uncover a novel physiological role for the *C. elegans* mitochondrial sirtuins, *sir-2.2* and *sir-2.3,* in lifespan regulation. Using a genetic approach, we demonstrate that *sir-2.2* and *sir-2.3* mutants live 28-30% longer than controls when fed the normal lab diet of *E. coli* OP50. Interestingly, this effect is diet specific and is not observed when animals are fed the strain HT115, which is typically used for RNAi experiments. While decreased consumption of food is a known mechanism for lifespan extension, this does not account for the increased lifespan in the mitochondrial sirtuin mutants. *sir-2.2* and *sir-2.3* mutants display altered expression of genes involved in oxidative stress response, including increased expression of the mitochondrial superoxide dismutase *sod-3* and decreased levels of catalases *ctl-1* and *ctl-2*. Like their extended lifespan phenotype, these alterations in oxidative stress gene expression are diet dependent. The mitochondrial sirtuin mutants are more resistant to the lifespan extending effects of low levels of superoxide, suggesting that their increased lifespan involves a hormetic response. Our data suggest that *sir-2.2* and *sir-2.3* are not completely redundant in function and may possess overlapping yet distinct mechanisms for regulating oxidative stress response and lifespan.

## Introduction

Sirtuins are a highly conserved family of NAD^+^-dependent enzymes that use NAD^+^ as a cofactor to execute mono-ADP ribosylation, deacetylation and a variety of other deacylation reactions (Blander & Guarente, 2004; Haigis et al., 2006; Houtkooper, Pirinen, & Auwerx, 2012). Distinct sirtuin family members are active in the cytoplasm, the nucleus, and the mitochondria, and act as molecular sensors and regulators of the cell’s metabolic state (Guarente, 2011; Haigis & Sinclair, 2010; He, Newman, Wang, Ho, & Verdin, 2012). Sirtuins are named for the *Saccharomyces cerevisiae* protein Sir2 (<u>s</u>ilent information <u>r</u>egulator 2), which increases yeast replicative lifespan when overexpressed (Kaeberlein, McVey, & Guarente, 1999). Genes and proteins with roles in lifespan extension have attracted much attention. As such, Sir2 is a well-studied protein, and its orthologs have also been extensively explored in many model systems. For example, over-expression of the *C. elegans* nuclear sirtuin SIR-2.1 was first shown to share lifespan enhancing functions with Sir2 more than a decade ago (Tissenbaum & Guarente, 2001). More recently, the robustness and extent of the lifespan functions of SIR-2.1 have been questioned (Burnett et al., 2011). Nonetheless extensive research on SIR-2.1 in *C. elegans* and orthologous sirtuins in other systems supports the model that the protein is a key player in modulating progression of aging as well as healthspan phenotypes via regulation of metabolism and oxidative stress responses (Aka, Kim, & Yang, 2011; Chang & Guarente, 2014; Guarente, 2013).

Mammals have seven sirtuins, SIRT1-7. Three of these sirtuins, SIRT3-5, reside in the mitochondria (Houtkooper et al., 2012). SIRT3 is a deacetylase that regulates the activity of various metabolic enzymes and the mitochondrial manganese-dependent superoxide dismutase SOD2 (Qiu, Brown, Hirschey, Verdin, & Chen, 2010; Schwer, Bunkenborg, Verdin, Andersen, & Verdin, 2006; Tao et al., 2010). It is the major deacetylase in the mitochondria (Lombard et al., 2007). The enzymatic functions of SIRT4 are the least characterized of the mitochondrial sirtuins (Haigis et al., 2006; He et al., 2012). SIRT4 is an ADP-ribosyltransferase that inhibits glutamate dehydrogenase and regulates insulin secretion (Ahuja et al., 2007; Haigis et al., 2006). Recently, studies have revealed the ability of SIRT4 to deacetylate malonyl CoA decarboxylase and control lipid homeostasis (Laurent et al., 2013). This is the first reported deacetylase activity for SIRT4, which has long thought to lack deacetylase activity (Ahuja et al., 2007). SIRT5 possesses demalonylase and succinylase activities (Du et al., 2011) and modifies the carbamoyl phosphate synthetase CPS1 to regulate the urea cycle (Du et al., 2011; Nakagawa, Lomb, Haigis, & Guarente, 2009).

In *C. elegans*, there are four sirtuins, SIR-2.1 to SIR-2.4. Two, SIR-2.2 and SIR-2.3, localize in the mitochondria (Wirth et al., 2013). While SIR-2.1 is well-studied, there is much less known about the biological roles of SIR-2.2 and SIR-2.3. These mitochondrial sirtuin genes are located adjacent to each other on chromosome X and share 75.3% sequence identity, suggesting that one developed from a gene duplication event (Wirth et al., 2013). They are orthologs of the mammalian SIRT4 protein (Jedrusik-Bode, 2014; Wirth et al., 2013). SIR-2.2 and SIR-2.3 in *C. elegans* physically interact with the mitochondrial biotin-dependent enzymes pyruvate carboxylase, propionyl-CoA carboxylase, and methylcrotonyl-CoA carboxylase (Wirth et al., 2013). Yet, the biological and physiological functions of SIR-2.2 and SIR-2.3 in *C. elegans* are largely unknown, and the mammalian SIRT4 protein is not fully studied (Haigis et al., 2006). We have explored the biological function of mitochondrial SIRT4 proteins using a genetic approach in *C. elegans*. We reveal that mutation of either of the *C. elegans* mitochondrial sirtuins *sir-2.2* and *sir-2.3* results in an extended lifespan, revealing novel roles in lifespan regulation and lack of redundancy of these similar proteins.

## Results

### Mitochondrial sirtuin mutants live longer than N2 when fed *E. coli* OP50 *ad libitum*

We acquired *C. elegans* sirtuin mutant strains to determine if any sirtuin plays a role in mediating phenotypic outcomes in a variety of mutants with defects in biosynthesis of NAD^+^, the sirtuin co-substrate. *sir-2.2(tm2673)* and *sir-2.3(ok444)* are deletion alleles that each remove portions of the sirtuin domain (see Materials and Methods, Wirth et al. 2013). In the process of examining the mutant strains over a developmental time course, we noted that the mitochondrial sirtuin mutants were more robust than the N2 control strain as they approached old age. We directly examined the aging of the animals by comparing the lifespan of the *sir-2.2(tm2673)* and *sir-2.3(ok444)* mutants relative to the control N2 strain to which they were backcrossed three times. Loss of *sir-2.2* or *sir-2.3* function resulted in an unexpected 28-30% increase in lifespan compared to the control N2 strain when fed *E. coli* OP50 *ad libitum* (Figure 1a,b). *sir-2.2* mutants lived an average of 4.2 days longer than N2, and *sir-2.3* mutants lived an average of 3.9 days longer than N2. This increased lifespan was consistently observed across four to five independent experiments (Table 1). To test whether the increased lifespan observed in single mitochondrial sirtuin mutants was due to upregulation of the remaining mitochondrial sirtuin, we measured mRNA levels of *sir-2.3* in the *sir-2.2(tm2673)* mutant and mRNA levels of *sir-2.2* in the *sir-2.3(ok444)* mutant. No compensatory upregulation was observed (Figure 1c).

**Figure 1.**
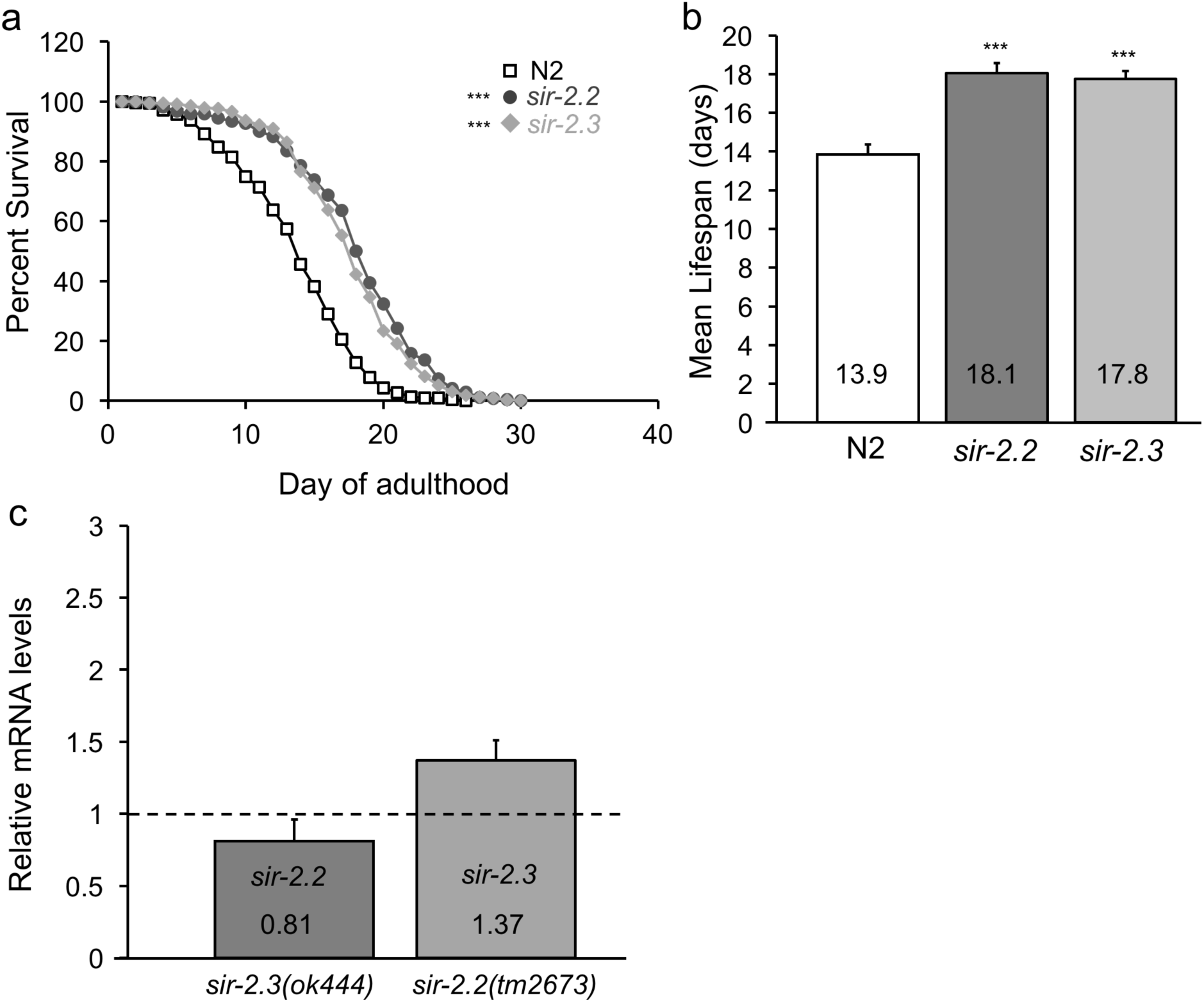
*sir-2.2(tm2673)* and *sir-2.3(ok444)* lifespan on OP50. (a) Survival curve of N2 control, *sir-2.2(tm2673)* and *sir-2.3(ok444)* fed *ad libitum* on OP50 across 4-5 experiments (See Table 1), ***p< 0.001, log rank t-test. **(b)** Mean lifespan values ±SEM of N2 control, *sir-2.2(tm2673)* and *sir-2.3(ok444)* when animals were fed OP50. Statistical analysis performed via one-way ANOVA followed by Tukey’s post hoc test, where ***p< 0.001. **(c)** Relative mRNA levels of *sir-2.2* in *sir-2.3(ok444)* and *sir-2.3* in *sir-2.2(tm2673)* measured with qRT-PCR. Values are the average of three independent experiments, each done in triplicate across two biological replicates, ±SEM

**Table 1.** Mean lifespan values of N2, *sir-2.2(tm2673)*, and *sir-2.3(ok444)* fed *ad libitum* on OP50

| Experiment | Genotype | Mean Lifespan±SEM | n | p-value (one-way ANOVA) |
| --- | --- | --- | --- | --- |
| 1 | N2 | 15.92 ± 0.68 | 41 | p=0.017840 |
| | *sir-2.3(ok444)* | 17.83 ± 0.56 | 57 | |
| 2 | N2 | 10.54 ± 0.67 | 64 | p<0.0001 |
| | *sir-2.2(tm2673)* | 19.98 ± 0.75 | 67 | |
| | *sir-2.3(ok444)* | 19.87 ± 0.59 | 64 | |
| 3 | N2 | 14.27 ± 0.48 | 64 | p<0.0001 |
| | *sir-2.2(tm2673)* | 17.71 ± 0.67 | 62 | |
| | *sir-2.3(ok444)* | 17.52 ± 0.66 | 60 | |
| 4 | N2 | 13.46 ± 0.53 | 64 | p<0.0001 |
| | *sir-2.2(tm2673)* | 16.22 ± 0.51 | 74 | |
| | *sir-2.3(ok444)* | 16.57 ± 0.51 | 71 | |
| 5 | N2 | 15.07 ± 0.31 | 74 | p<0.0001 |
| | *sir-2.2(tm2673)* | 18.30 ± 0.34 | 69 | |
| | *sir-2.3(ok444)* | 17.02 ± 0.32 | 77 | |

### Lifespan extension due to loss of *sir-2.2* or *sir-2.3* is diet dependent

In control experiments in preparation for RNAi experiments, we noted that food source had an effect on lifespan of *sir-2.2* and *sir-2.3* mutants. Feeding with the *E. coli* HT115 strain used for administering RNAi had no effect on lifespan of N2 control animals but decreased the lifespan of *sir-2.2* and *sir-2.3* mutants (Figure 2a,b,c). The lifespan of *sir-2.2(tm2673)* on HT115 is 73% of its lifespan on OP50, or around four days shorter when fed HT115 compared to OP50 (Figure 2b,c). The lifespan of *sir-2.3(ok444)* on HT115 is 77% of its lifespan on OP50, or around three days shorter when fed HT115 compared to OP50 (Figure 2b,c). Interestingly, the survival and mean lifespan of *sir-2.3(ok444)* was still significantly extended compared to N2 when fed HT115 whereas that of *sir-2.2(tm2673)* on HT115 was no different than N2 (Figure 2a,b,c). Therefore, growth on HT115 appears to eliminate the lifespan extension of *sir-2.2(tm2673)* relative to N2 and reduces that of *sir-2.3(ok444)*, suggesting that the extended lifespan phenotypes of *sir-2.2(tm2673)* and *sir-2.3(ok444)* are both diet dependent, but the diet effect is more pronounced in *sir-2.2(tm2673)* relative to *sir-2.3(ok444).*

**Figure 2.**
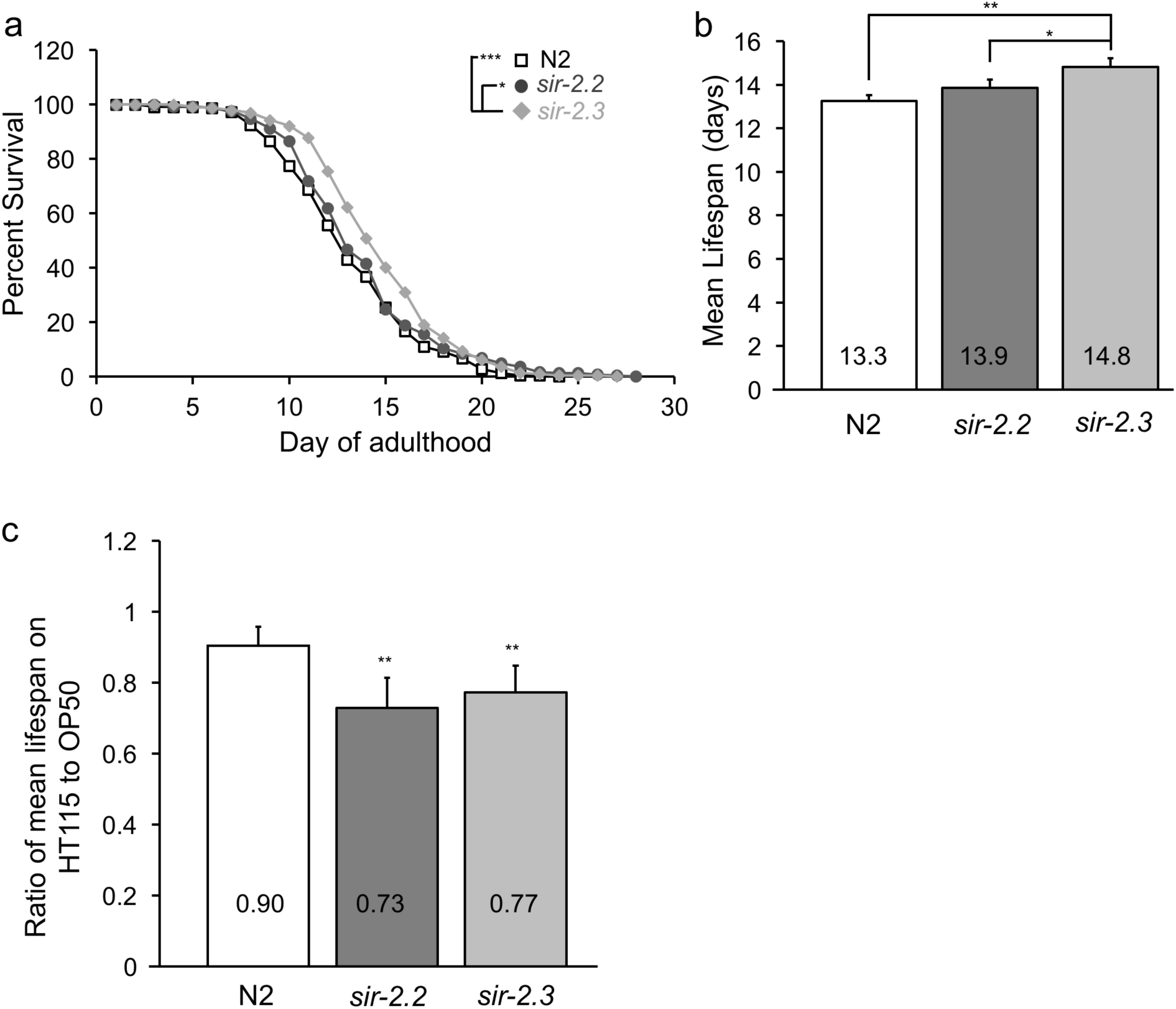
*sir-2.2(tm2673)* and *sir-2.3(ok444)* lifespan on HT115. (a) Survival curve of N2 control, *sir-2.2(tm2673)*, and *sir-2.3(ok444)* animals fed *ad libitum* on HT115 across four experiments (See Table S1). *sir-2.3(ok444)* has increased survival compared to N2 (***p< 0.001) and to *sir-2.2(tm2673)* (*p< 0.05) on HT115, log rank t-test. **(b)** Mean lifespan values ±SEM of N2 control, *sir-2.2(tm2673)* and *sir-2.3(ok444)* when animals were fed HT115. Statistical analysis performed via one-way ANOVA followed by Tukey’s post hoc test, where **p< 0.01, *p< 0.05. **(c)** Ratios of mean lifespan values of animals on HT115 to mean lifespan values of animals on OP50 (See Figure 1b). Statistical analysis performed via one sample t-test comparing to 1 where **p< 0.01.

### Changes in food intake do not contribute to the extended lifespan of *sir-2.2* and *sir-2.3* mutants

Because dietary restriction increases lifespan (Sohal & Weindruch, 1996), we investigated whether the lifespan extension of the mitochondrial sirtuin mutants was due to decreased food intake. Muscular contractions of the pharynx pump food into the *C. elegans* intestine (Avery & You, 2012). Thus, we measured pharyngeal pumping rates as an indicator of food intake. We measured the pharyngeal pumping rate of N2, *sir-2.2(tm2673)*, and *sir-2.3(ok444)* fed OP50 *ad libitum* or fed OP50 for five minutes after a six hour fasting period. There was no observed difference between the N2 control and the mitochondrial sirtuin mutants whether they were fed *ad libitum* or post-fasting, suggesting that decreased food intake does not contribute to the increase in lifespan of *sir-2.2(tm2673)* or *sir-2.3(ok444)* (Figure 3a,b).

**Figure 3.**
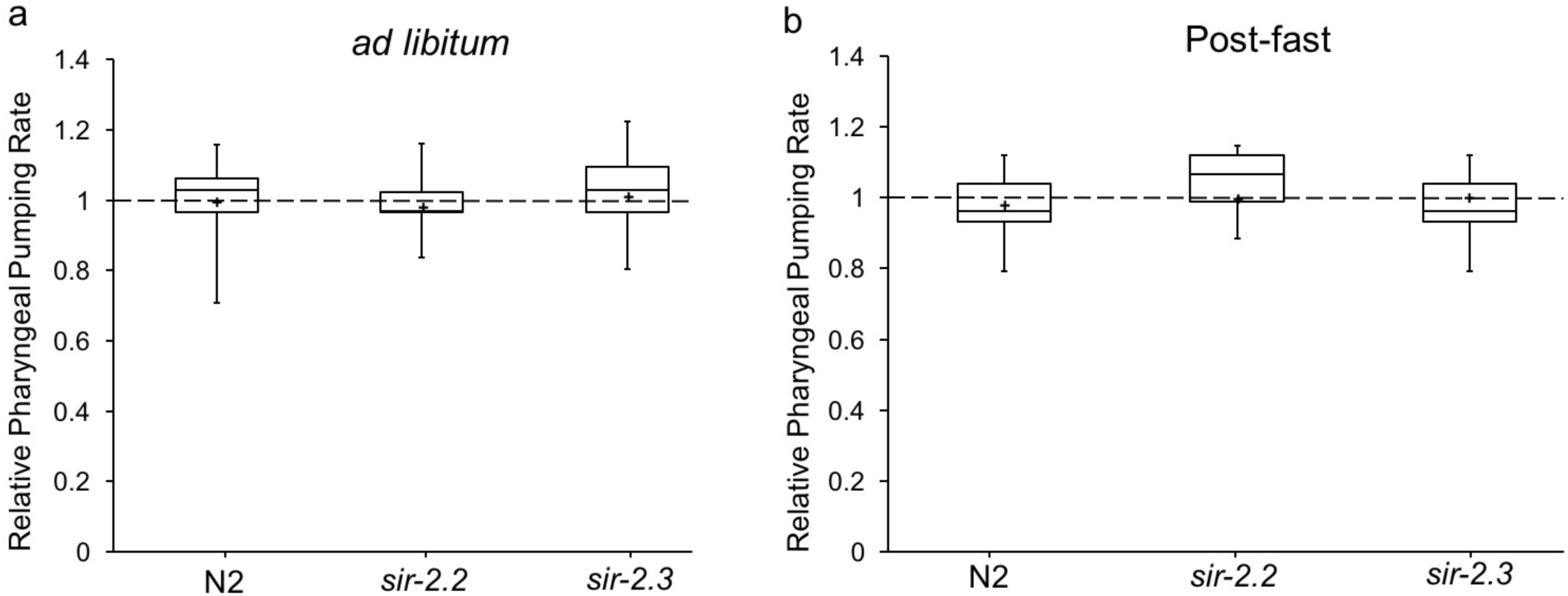
Relative pharyngeal pumping rates of *sir-2.2(tm2673)* and *sir-2.3(ok444)*. (a) Animals fed OP50 *ad libitum*, N2 (n=30), *sir-2.2(tm2673)* (n=33), *sir-2.3(ok444)* (n=34). **(b)** Animals fed OP50 for 5 minutes after six hour fasting period, N2 (n=28), *sir-2.2(tm2673)* (n=33), *sir-2.3(ok444)* (n=36). Values are from three independent experiments. Box plot displays minimum and maximum values, first quartile, median, and third quartile, where + denotes the mean value.

### A hormetic response may underlie the lifespan extension produced by loss of the mitochondrial sirtuins

Both *sir-2.2(tm2673)* and *sir-2.3(ok444)* mutants are hypersensitive to high levels of oxidative stress (Wirth et al., 2013), suggesting that the proteins might help ameliorate stress and additionally may experience an elevated constitutive level of oxidative stress. We hypothesized that a mounted response to constitutive mild oxidative stress potentially experienced by *sir-2.2* and *sir-2.3* mutants, a hormetic response, could mediate the extended lifespan. To investigate this hypothesis, we first examined the mutant strains for evidence of a stress response by measuring the mRNA levels of key oxidative stress response genes including the mitochondrial superoxide dismutases *sod-2* and *sod-3*, the cytoplasmic catalase *ctl-1*, and the peroxisomal catalase *ctl-2*. *sod-2* mRNA levels were unchanged in either mitochondrial sirtuin mutant (Figure S1). *sod-3* mRNA levels were elevated more than three-fold in *sir-2.2* mutants and more than two-fold in *sir-2.3* (Figure 4a). In contrast, levels of *ctl-1* and *ctl-2* messages were depressed approximately two-fold in both *sir-2.2* and *sir-2.3* compared to N2 (Figure 4a).

**Figure 4.**
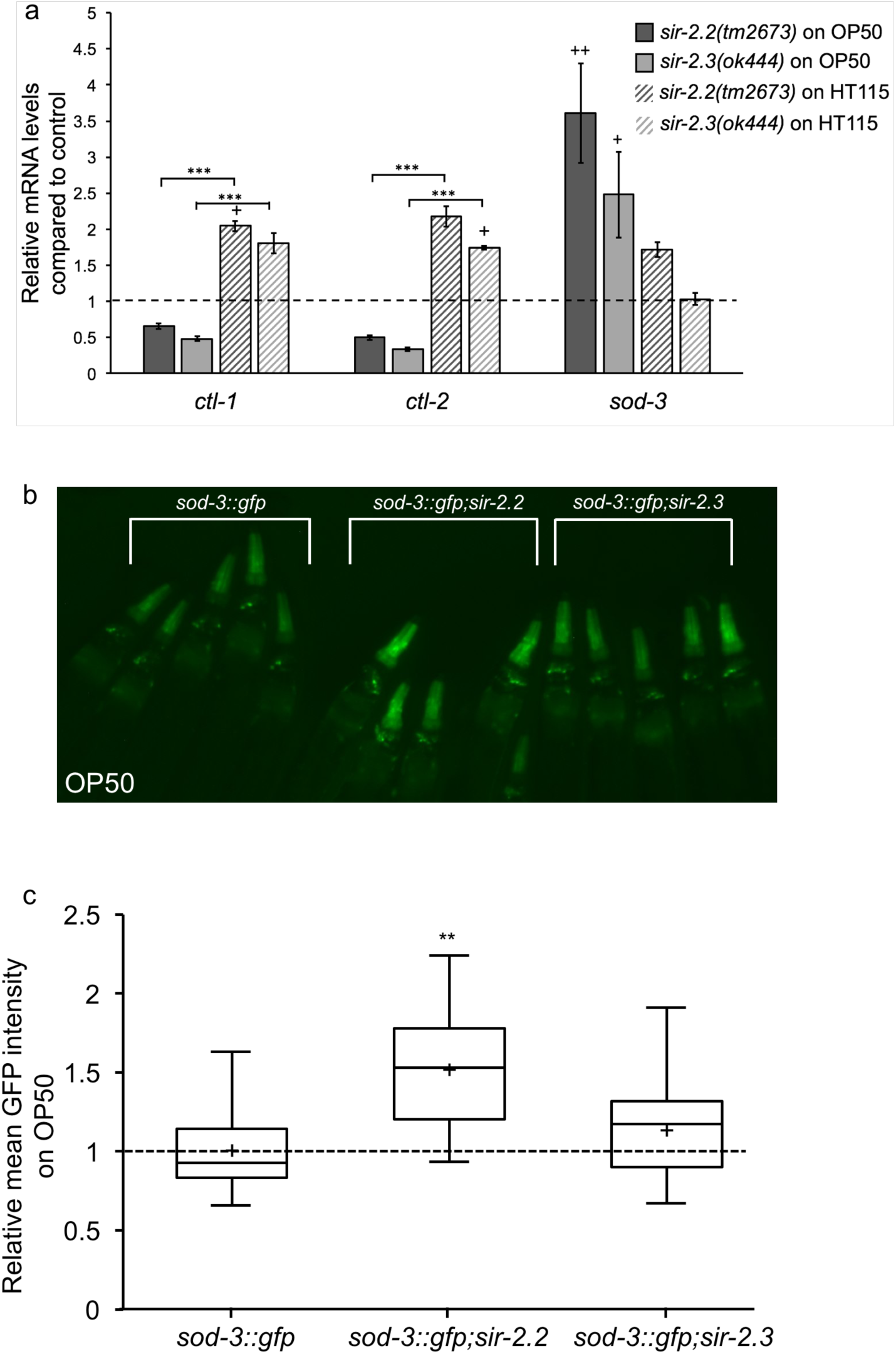
*sod-3, ctl-1,* and *ctl-2* expression in *sir-2.2(tm2673)* and *sir-2.3(ok444)* on OP50. (a) qRT-PCR was used to measure relative mRNA levels of *ctl-1, ctl-2,* and *sod-3* in N2 control, *sir-2.2(tm2673)*, and *sir-2.3(ok444)* fed either OP50 or HT115. Values are averages from two biological replicates, both done in triplicate, error bars represent SEM, ***p< 0.001 using Student’s t-test, +p< 0.05, ++p< 0.01 using one sample t-test comparing to 1. **(b)** Image representative of the relative GFP levels in the pharynx of *psod-3::gfp, psod-3::gfp;sir-2.2*, and *psod-3::gfp;sir-2.3* animals fed OP50. Image is of five day 1 adult animals of each genotype placed adjacent to one another, paralyzed with 1 mM levamisole and imaged simultaneously. **(c)** Quantification of the mean GFP intensity in the pharynx of *sod-3::gfp*, *sod-3::gfp;sir-2.2*, and *sod-3::gfp;sir-2.3* animals on OP50. Sample sizes were 33 to 35. Box plot displays minimum and maximum values, first quartile, median, and third quartile, where + denotes the mean value. Statistical analysis performed via one-way ANOVA followed by Tukey’s post hoc test where **p< 0.01.

To test whether food source affects regulation of *sod-3*, *ctl-1,* and *ctl-2* in the mitochondrial sirtuin mutants, we measured the mRNA levels of these genes in control and mutant animals fed HT115. On HT115, *sir-2.2* and *sir-2.3* mutants show approximately a two-fold increase in *ctl-1* and *ctl-2* mRNA levels as opposed to a two-fold decrease when these animals are fed OP50 (Figure 4a). Unlike the two to three-fold increase in *sod-3* in the mitochondrial sirtuin mutants fed OP50, *sod-3* expression is unchanged compared to the control strain (Figure 4a). Thus, the detected gene expression changes correlate with lifespan. These data support the hypothesis that the changes in *sod-3*, *ctl-1*, and *ctl-2* expression observed when mitochondrial sirtuin mutant animals are fed OP50 may play a role in the extended lifespan of *sir-2.2(tm2673)* and *sir-2.3(ok444)*.

To seek independent evidence for the changes in *sod-3* expression in *sir-2.2* and *sir-2.3*, we placed a *psod-3::gfp* transgene in both the *sir-2.2* and *sir-2.3* mutant background and measured *sod-3* expression levels via intensity of GFP expression in the pharynx. On OP50, there was a significant increase in mean GFP intensity in *sir-2.2* mutants whereas the mean GFP intensity of *sod-3*::GFP was slightly increased in *sir-2.3* mutants. This increase did not rise to the level of statistical significance (Figure 4b,c).

To further test the hypothesis that a hormetic response contributes to the extended lifespan of the mitochondrial sirtuin mutants, we treated the animals with low or high levels of paraquat, a superoxide radical generator. Low levels of superoxide generators, such as 0.1 mM of paraquat, extend the lifespan of wild-type worms (Yang & Hekimi, 2010). If the mitochondrial sirtuin mutants are already experiencing elevated oxidative stress, we suspected that 0.1 mM of paraquat would not extend their lifespans further. Indeed N2 worms treated with 0.1 mM paraquat lived 35.3% longer, extending their lifespan to 18.8 days from 13.9 days (Figure 5a,e). *sir-2.2* mutants did not show an extended lifespan whereas *sir-2.3* mutants lived approximately 17.6% longer, extending their lifespan to 20 days from 17 days (Figure 5b,c,e). This supports the hypothesis that a hormetic response in both *sir-2.2* and *sir-2.3* mutants mediates their increased lifespan. Because *sir-2.3* mutant lifespan was increased upon treating with low levels of paraquat, a hormetic response may be mediating the lifespan extension in the *sir-2.3* mutant to a lesser degree than *sir-2.2* mutants. As previously published, *sir-2.2* and *sir-2.3* mutants were more sensitive than controls to the high concentration of 3.75 mM paraquat (Figure 5d,e, Wirth et al. 2013).

**Figure 5.**
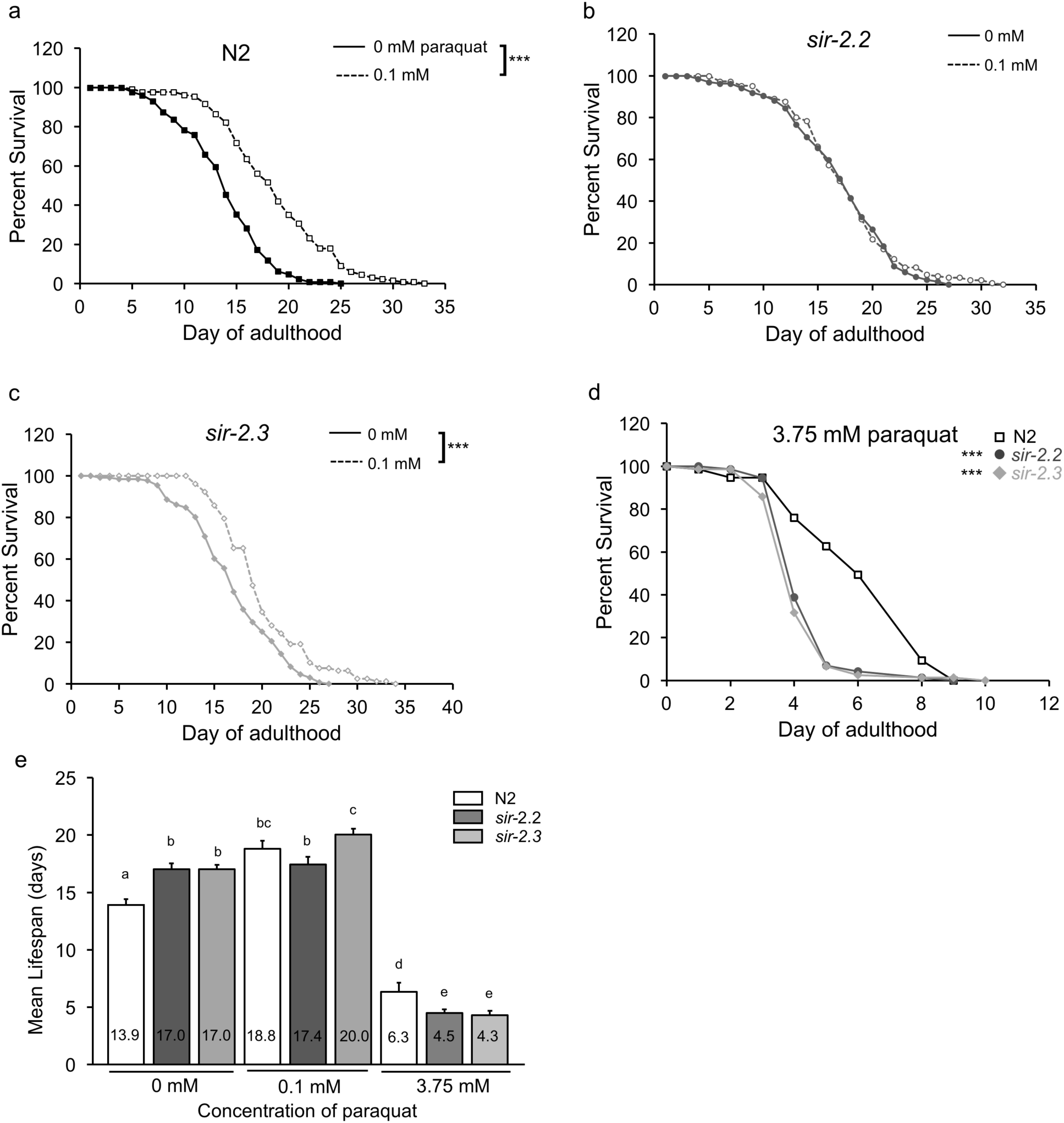
*sir-2.2(tm2673)* and *sir-2.3(ok444)* lifespan on varying concentrations of paraquat. (a,b,c) Survival curve of N2, *sir-2.2(tm2673)*, and *sir-2.3(ok444)* treated with 0.1 mM paraquat and untreated, ***p< 0.001, log rank t-test. **(d)** Survival curve of N2, *sir-2.2(tm2673)*, and *sir-2.3(ok444)* treated with 3.75 mM paraquat, ***p< 0.001, log rank t-test. **(e)** Mean lifespan values ±SEM of N2, *sir-2.2(tm2673)*, *sir-2.3(ok444)* fed OP50 and treated with 0.1 mM, 1.0 mM, 3.75 mM or no paraquat. Values are from three independent experiments (see Table S2), one-way ANOVA followed by Tukey’s post hoc test where different letters represent a statistical difference of p< 0.01.

## Discussion

In this study, we uncovered a novel role for the mitochondrial sirtuins *sir-2.2* and *sir-2.3* in lifespan regulation; removing either of their activities increases lifespan by more than 25 %. Interestingly, *sir-2.1* mutants have also been reported to have an increased lifespan compared to controls when fed OP50 *ad libitum* (Moroz et al., 2014). Here, we reveal that the lifespan extending mechanisms in *sir-2.2* and *sir-2.3* mutants are diet dependent and mediated in part by response to oxidative stress. Expression analyses have indicated that SIR-2.2 and SIR-2.3 are not functionally redundant as they are expressed at different times during embryogenesis and have shown to be localized in different tissues (Wirth et al., 2013). Our results support their non-redundancy and uncover the presence of stress-related phenotypic differences between the two.

When fed the *E. coli* strain HT115, *sir-2.2* and *sir-2.3* mutants no longer had the increased lifespan present on OP50. However, *sir-2.3* mutants still had an extended lifespan compared to N2 when fed HT115. The lifespan of *C. elegans* is regulated by their ability to respond to changes in diet (Pang & Curran, 2014). When fed either OP50 or HT115, N2 animals have similar lifespans, as observed in this study and published elsewhere (Brooks, Liang, & Watts, 2009). This ability to adapt to different diets seems to involve *sir-2.2* and *sir-2.3*, as *sir-2.2* and *sir-2.3* mutant worms do not have the same lifespan upon a switch to a different food source, unlike control animals. It is necessary to investigate the mechanism used by *C. elegans* in response to changes in diet to uncover the specific role mitochondrial sirtuins may play in this process.

Curiously, unlike *age-1* and *daf-2* mutants, which have increased catalase activity (Larsen, 1993; Vanfleteren, 1993), both *sir-2.2* and *sir-2.3* exhibit an approximate two-fold decrease in catalase mRNA levels, an unexpected result due to catalase’s function in detoxifying hydrogen peroxide. Although cytosolic catalase is required to extend lifespan of *daf-2* mutants, catalase inactivation extended the lifespan of *Saccharomyces cerevisiae* due to induction of superoxide dismutase (Mesquita et al., 2010). It is not well understood why both *sir-2.2* and *sir-2.3* may decrease their catalase levels.

While the lifespan extending mechanism in *sir-2.2* and *sir-2.3* mutants still requires investigation, our works emphasizes the importance of the sirtuin family as modulators of oxidative stress response and lifespan and broadens our ability to target this class of proteins for new therapies, potentially in ways that can improve healthspan.

## Materials and Methods

### Nematode strains and maintenance

*C. elegans* strains were maintained using standard methods at 20° C on *E. coli* OP50 (Brenner, 1974). We used the strains N2 Bristol as the wild-type control, RB654 *sir-2.3(ok444)*, and CF1553 *muIs84*[(*pAD476*) *sod-3p::*GFP + *rol-6*] which were provided by *Caenorhabditis* Genetics Center (CGC). We also used *sir-2.2(tm2673),* which was provided by the Mitani lab through the National Bio-Resource Project of the MEXT, Japan. *sir-2.2(tm2673)* and *sir-2.3(ok444)* were each outcrossed three times into the N2 strain that was used for a control for all experiments.

*sir-2.2(tm2673)* is a deletion allele that removes exons 3 and 4, corresponding to 75 amino acids of the sirtuin domain (Wirth et al., 2013). The deletion allele *sir-2.3(ok444)* removes part of exon 3, all of 4 and 5 and a portion of exon 6, generating an in-frame stop codon, resulting in a truncation of 141 amino acids from the C terminus of the protein (Wirth et al., 2013).

### Lifespan Analysis

Lifespan assays were conducted at 20°C on standard NGM plates with 400 µl of *E. coli* OP50 or HT115 and were replicated in at least three independent experiments. Animals were synchronized using a timed egg lay or an egg preparation (Sulston & Hodgkin, 1988). 30 L4 animals were placed on individual plates at the start of the assay and moved to new plates every day. To assess survival, worms were prodded with a platinum wire every day and scored as dead if non-responsive. Worms with internal hatching or an “exploded” phenotype were censored.

For paraquat lifespan analysis, 200 µl of paraquat (methyl viologen dichloride hydrate 98%, Sigma) diluted in water was added to NGM plates spotted with 400 µl OP50 for a final concentration of 3.75 mM, 1.0 mM or 0.1 mM paraquat. 30 synchronized L4 worms were placed on individual plates at the start of the assay and transferred to new plates every day. Results represent an average of three independent experiments.

### Pharyngeal Pumping

Pharyngeal pumping was measured in day one adults fed *ad libitum* or five minutes post six hour fasting period on OP50 as described (Lemieux et al., 2015). All worms were grown on OP50 at 20° C. Pumping rates were measured in ten second intervals using a Nikon SMZ1500 stereoscope equipped with Roper Scientific Photometrics CoolSnap EZ camera.

### Quantitative RT-PCR

RNA was isolated from mixed stage worms using TRIZOL reagent (Invitrogen). 1 µg of RNA was converted to cDNA using the qScript cDNA Synthesis Kit (Quanta Biosciences). cDNA was diluted 1:10 and used for quantitative PCR using SYBR Green and Applied Biosciences RT-PCR machine. A combination of three control primer sets (*cdc-42*, *tba-1, and pmp-3*) were used. *cdc-42* F: ctgctggacaggaagattacg; R: ctgggacattctcgaatgaag

*tba-1* F: gtacactccactgatctctgctgaca; R: ctctgtacaagaggcaaacagccatg

*pmp-3* F: gttcccgtgttcatcactcat; R: acaccgtcgagaagctgtaga

*sir-2.2* F: ggtatcccagattaccgctcg; R: ccaaatctcggccaggctaa

*sir-2.3* F: ggaacttcatggcaacgctc; R: gaacccttgttcgctaccca

*sod-2* F: gaggcggtctccaaaggaaa; R: ccagagatccgaagtcgctc

*sod-3* F: ctccaagcacactctcccag; R: tccctttcgaaacagcctcg

*ctl-1* F: acaaggagacgtatccaaaacc; R: tccagcgaccgttgaaaaac

*ctl-2* F: ctacagtcggtggtgagagc; R: tacccatctgggagtcctcg

Results represent the average of two independent biological samples unless otherwise denoted, each of which was amplified in triplicate.

### sod-3::GFP expression quantification

Day 1 adult animals were placed on unspotted NGM plates, treated with 1 mM levamisole to restrict movement, and imaged on a Nikon SMZ1500 stereoscope. Images were collected using Roper Scientific Photometrics CoolSnap EZ camera using a 19.05 second exposure at 10x magnification. Images were analyzed using ImageJ to measure mean GFP intensity in the pharynx of each animal with background removed.

## Acknowledgements

Strains were provided by the Caenorhabditis Genetics Center, which is funded by NIH Office of Research Infrastructure Programs (P40 OD010440), and the Mitani Lab through the National Bio-Resource Project of the MEXT, Japan. This work was supported by National Institutes of 286 Health award number GM086786 to WHR.

## Contributions

SMC conceived and coordinated the study. SMC performed experiments in Figures 1 through 5 and analyzed results. SMC and MRM performed experiments in Figure 4a and SMC analyzed results. SMC and WHR wrote the paper.

## Chang et al, Supplementary Tables and Figures

**Table S1.** Mean lifespan values of N2, *sir-2.2(tm2673)*, and *sir-2.3(ok444)* fed *ad libitum* on HT115

| Experiment | Genotype | Mean Lifespan±SEM | n | p-value (one-way ANOVA) |
| --- | --- | --- | --- | --- |
| 1 | N2<br>*sir-2.2(tm2673)*<br>*sir-2.3(ok444)* | 14.21±0.51<br>16.00±0.61<br>16.37±0.42 | 71<br>65<br>70 | p=0.009 |
| 2 | N2<br>*sir-2.2(tm2673)*<br>*sir-2.3(ok444)* | 12.57±0.43<br>12.77±0.34<br>12.59±0.31 | 47<br>48<br>43 | p=0.896 |
| 3 | N2<br>*sir-2.2(tm2673)*<br>*sir-2.3(ok444)* | 13.31±0.43<br>13.26±0.43<br>14.84±0.60 | 62<br>68<br>51 | p=.037 |
| 4 | N2<br>*sir-2.2(tm2673)*<br>*sir-2.3(ok444)* | 12.37±0.38<br>13.35±0.35<br>14.55±0.30 | 49<br>70<br>63 | p=.000179 |

**Table S2.**
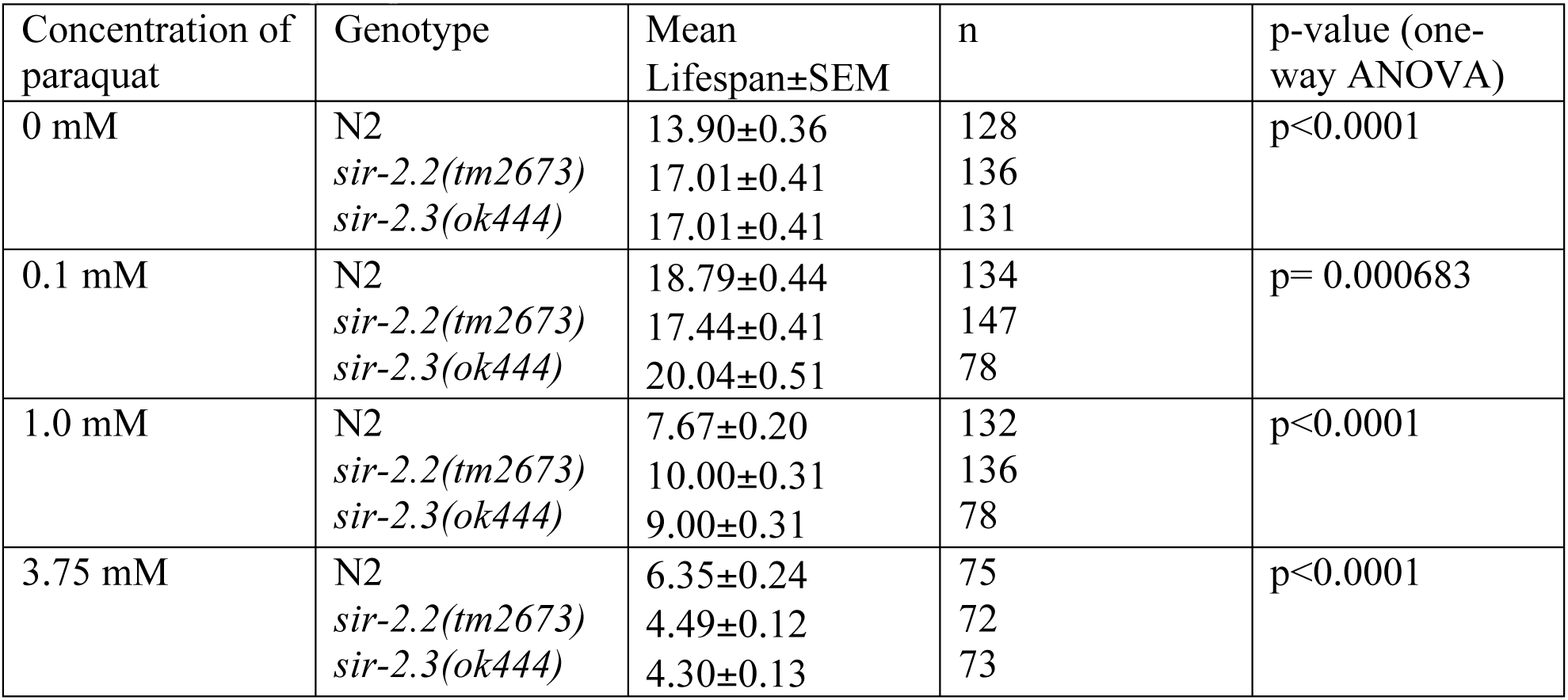
Mean lifespan values of N2, *sir-2.2(tm2673)*, and *sir-2.3(ok444)* treated with varying concentrations of paraquat

| Concentration of paraquat | Genotype | Mean Lifespan±SEM | n | p-value (one-way ANOVA) |
| --- | --- | --- | --- | --- |
| 0 mM | N2<br>*sir-2.2(tm2673)*<br>*sir-2.3(ok444)* | 13.90±0.36<br>17.01±0.41<br>17.01±0.41 | 128<br>136<br>131 | p<0.0001 |
| 0.1 mM | N2<br>*sir-2.2(tm2673)*<br>*sir-2.3(ok444)* | 18.79±0.44<br>17.44±0.41<br>20.04±0.51 | 134<br>147<br>78 | p= 0.000683 |
| 1.0 mM | N2<br>*sir-2.2(tm2673)*<br>*sir-2.3(ok444)* | 7.67±0.20<br>10.00±0.31<br>9.00±0.31 | 132<br>136<br>78 | p<0.0001 |
| 3.75 mM | N2<br>*sir-2.2(tm2673)*<br>*sir-2.3(ok444)* | 6.35±0.24<br>4.49±0.12<br>4.30±0.13 | 75<br>72<br>73 | p<0.0001 |

**Figure S1.**
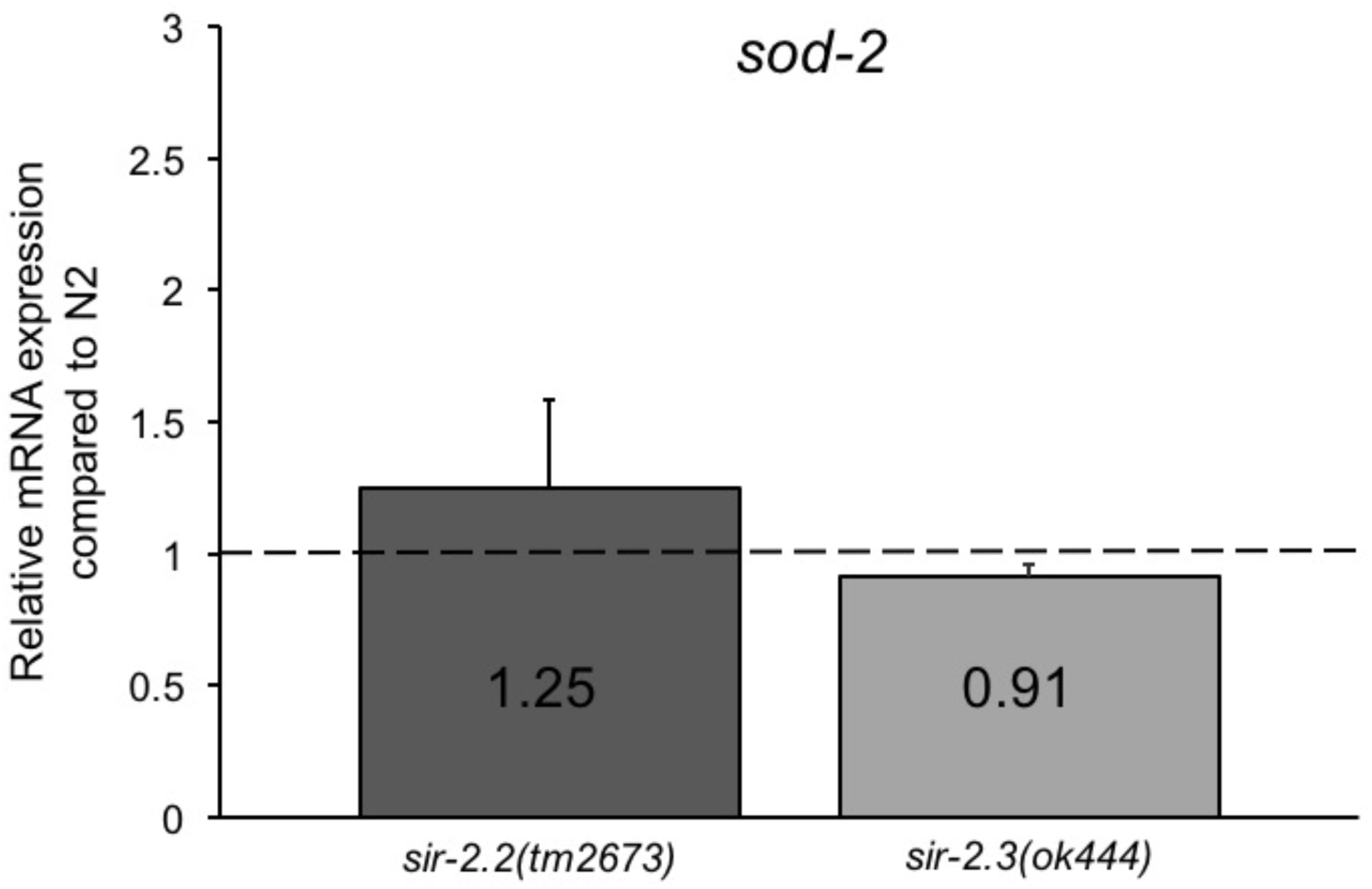
*sod-2* mRNA levels in mitochondrial sirtuin mutants *sir-2.2(tm2673)* or *sir-2.3(ok444)*. mRNA expression was measured with qRT-PCR and compared to N2 control. Values are averages from two biological replicates, each done in triplicate, error bars represent SEM.

